# ZSeekerDB: A database of Z-nucleic acid sequences across organismal genomes

**DOI:** 10.64898/2025.12.24.696366

**Authors:** Spyros Zaranikas, Georgios Megalovasilis, Kimonas Provatas, Guliang Wang, Apostolos Zaravinos, Karen Vasquez, Georgakopoulos-Soares

## Abstract

Alternative (non-B) nucleic acid structures such as Z-nucleic acids are emerging as key regulators of genome function. The density and distribution of Z-nucleic acid sequences across organismal and viral genomes can provide insights into their biological roles and evolutionary trajectory. Using our recently developed ZSeeker algorithm, we systematically analyzed over 280,000 organismal genome assemblies, identifying more than 850 million putative Z-forming loci. We also incorporated genomic coordinates, Z-score, and taxonomic metadata, enabling cross-species comparative and functional analyses. We introduce ZSeekerDB, the first large-scale, multi-taxon database cataloging Z-nucleic acid sequences, across organisms representing all major branches of life. ZSeekerDB enables interactive searches, visualizations, and downloads of Z-nucleic acid sequence data for independent analysis. ZSeekerDB is implemented as a web-portal for browsing, analyzing and downloading Z-forming loci, publicly available at https://zseeker-db.com/.

## Introduction

The canonical right-handed DNA double helix, known as the B-DNA structure, was first described by Watson, Crick, Wilkins, and Franklin (1). However, DNA can also adopt alternative conformations, collectively referred to as non-B DNA (2). Z-DNA is a left-handed helical form of DNA characterized by a zig-zag backbone that typically forms in alternating purine-pyrimidine sequences under negative supercoiling stress and/or high-salt conditions (3). Since its discovery, Z-DNA has emerged as a dynamic structural element with roles in gene regulation, genome instability, and immune signaling (4–11). Double-stranded RNA can also adopt a similar left-handed helical conformation (Z-RNA) and has been implicated in antiviral surveillance and inflammation through its recognition by the Zα domain of ADAR1 and ZBP1 (12–14).

Accurate mapping of Z-DNA occupancy *in vivo* in living genomes presents significant technical challenges due to its transient and dynamic nature. Historically Z-DNA was detected biochemically using chemical modification footprints, Z-DNA-binding proteins, or antibodies (15, 16). In the human genome, Z-DNA was mapped using Zα pull-down and sequencing (17). However, most *in vivo* Z-DNA conformation detecting or mapping require pre-treatments or conditions that can interrupt the native Z-DNA formation. For example, the Zα domain is not only a probe, but also a strong Z-DNA formation inducer/stabilizer (18). In addition, the global chromatin structure, binding proteins, and DNA metabolism activities including replication and transcription could impact the access and binding of the protein or chemical probes to DNA, making the mapping more of a snapshot of accessible Z-DNA in the genome under a given condition.

The potential for DNA/RNA to adopt the Z-form is based on the fundamental chemical principles of sequence preference, making i*n silico* Z-DNA or Z-RNA prediction and mapping feasible. Bioinformatic detection of Z-DNA and Z-RNA generally relies on sequence-based predictors that recognize alternating purine-pyrimidine tracts, such as (CG)n or (CA/TG)n, which are known to have high Z-DNA-forming potential (19, 20). Tools such as Z-HUNT (21), DeepZ (22), Z-DNABERT (11) can systematically identify such loci. Recently, we published ZSeeker (23), a novel, experimentally guided and validated approach that we benchmarked against prior Z-DNA detection tools, enabling more precise identification of potential Z-nucleic acid sequences and addressing several limitations of existing software.

The rapid expansion of genome sequencing technologies has led to an unprecedented availability of genome assemblies across the tree of life. Advances in long-read sequencing, assembly algorithms, large-scale initiatives such as the Earth BioGenome Project, and various national biodiversity programs have made it possible to obtain high-quality assemblies for thousands of species, spanning all major domains of life and viral realms (24). This growing body of organismal genome assemblies provides a unique opportunity to study genome organization, evolutionary dynamics, and the distribution of sequence features across phylogenetically distant lineages, revealing both conserved and lineage-specific patterns of genomic architecture. Currently, the only database that hosts Z-nucleic acid sequences is non-B DB, which has genome-wide maps for twelve organisms (25). However, no databases are currently available that catalog Z-forming sequences across genomes spanning the tree of life.

Here, we present ZSeekerDB, a dedicated database for Z-nucleic acid sequences across genomes representing all major branches of the tree of life. The current release encompasses data from over 280,000 genomes, containing more than 850 million predicted Z-DNA- and Z-RNA-forming sequences. By integrating sequence-level predictions with rich taxonomic and assembly metadata, ZSeekerDB enables unparalleled cross-species comparisons of Z-nucleic acid density, composition, and evolutionary patterns. Thus, ZSeekerDB provides the first resource to provide Z-DNA/Z-RNA sequences at true phylogenomic scale.

## Materials and Methods

### Data collection

Complete genomic assemblies were downloaded from the NCBI GenBank and RefSeq repositories on 09-02-2025 (26, 27). Duplicate accessions were removed, choosing RefSeq over GenBank, as RefSeq entries are highly curated and quality-controlled by NIH. Each record was parsed to extract assembly descriptors including organism name, infraspecific name, taxonomy identifier, genome size and G+C content. Genome size was manually corrected with a bash script that removed -N stretches longer than 10 nucleotides from the total base count. We also manually included additional telomere-to-telomere (T2T) assemblies beyond those present in the above dataset, drawing from various sources besides NCBI (GenomeArk: https://github.com/genomeark/genomeark-update/, GigaScience Database: https://doi.org/10.5524/102657 (28), and Figshare repository: https://doi.org/10.6084/m9.figshare.25244542.v2 (29)). Our final dataset comprised 280,884 unique organismal genomes across all major taxonomic groups (**Supplementary Table 1**). The dataset was cross validated using NCBI’s Taxonomy database to confirm the consistency of organismal names and identifiers, before extracting full taxonomic lineages for each assembly. This multi-step filtering process ensured that the analyzed collection represented a comprehensive and high-quality corpus of organismal genome sequences suitable for large-scale comparative analysis of Z-DNA/Z-RNA-forming loci.

### Prediction of Z-DNA sequences

Z-DNA sequences were predicted across all genomes using the ZSeeker software (23) from https://github.com/Georgakopoulos-Soares-lab/ZSeeker. Jobs were launched via a custom SLURM script with default scoring and penalty parameters. ZSeeker produced a .csv file for each genome, with one prediction per row. Each prediction included columns for chromosome, start, end, sequence, and a numeric Z-DNA score.

### Database design and data pipelines

ZSeeker records were linked to assemblies via stable assembly identifiers. Supplementary metadata schema provided taxonomic descriptors (e.g., superkingdom, kingdom, genus, species name, NCBI taxid), assembly attributes (e.g., genome size, G+C content, assembly level) and Z-DNA/-RNA density metrics. SQL views materialized common joins so that sequence-level and species-level queries executed efficiently in DuckDB. The backend was implemented in Go, exposing a small set of read-only REST endpoints and a whitelisted SQL endpoint for parameterized SELECT queries used by the UI. The application accessed all data through a single DuckDB file. The front end was implemented in React (TypeScript) and rendered interactive visualizations using ECharts. Exports were streamed to the client to support large result sets.

### Web-interface

ZSeekerDB was built as a full-stack, web-based platform. The backend was implemented using the Gin Web Framework (v1.10), which provided a lightweight and high-performance foundation for scalable web services. Analytical processing and high-throughput querying of genomic datasets were supported by DuckDB, integrated via the go-duckdb driver (v1.8.3) to enable in-process OLAP functionality optimized for complex analytical workloads. The frontend, implemented in React (v19.1.0), delivered a responsive and interactive user environment. Material UI (v7.1.0) ensured a consistent and accessible design framework, while Apache ECharts (v5.6.0), integrated via ECharts-for-React (v3.0.2), enabled rich, high-resolution data visualizations. The application was built with Vite (v6.3.5) [MIT License] to support rapid development and optimized deployment and employed TypeScript (v5.8.3) [Apache License 2.0] to ensure type safety, code reliability, and maintainability. This architecture established a robust and high-performance computational environment tailored for real-time visualization and exploration of Z-DNA/Z-RNA data across genomic contexts.

### ZSeekerDB database overview and functionality

#### Database contents and usage

ZSeekerDB provides an interactive, web-based environment for the systematic exploration of Z-nucleic acid-forming sequences across over 280,000 complete organismal genome assemblies. The platform integrates multiple functional modules designed to facilitate both large-scale comparative analyses and fine-grained locus-level exploration. Users can browse assemblies through a faceted Species Browser, perform targeted Sequence Searches, and visualize global and chromosome-level trends through the Species Insights module. All analytical components operate directly on a unified DuckDB backend, ensuring high-performance queries and reproducible outputs.

Users visiting ZSeekerDB are navigated to the homepage (**Fig. 1A**) that displays general information about the database, with a navigation bar on the left providing access to all platform sections. The main sections include the Sequence Search, the Species Browser, and the Species Insights page, which provides visualizations. Additionally, the About, Help, and Privacy pages offer essential support and transparency to users of the ZSeekerDB platform. The About page outlines ZSeekerDB scope, underlying ZSeeker-based pipeline and data sources, species coverage, and links to publications and contact information. The Help page introduces the Z-DNA record concept and provides step-by-step guidance for core workflows including searching sequences, filtering by taxonomy, species, chromosome or score, exploring species-level views and visualizations, and exporting results. The Privacy page outlines the site’s data protection practices, security measures, and licensing terms, which adhere to the Creative Commons Attribution-NonCommercial-ShareAlike 4.0 (CC BY-NC-SA 4.0). Finally, the Download page offers downloadable, versioned datasets of predicted Z-DNA/-RNA loci in CSV, JSON, BED, and Parquet formats, supporting diverse bioinformatics queries.

**Figure 1:**
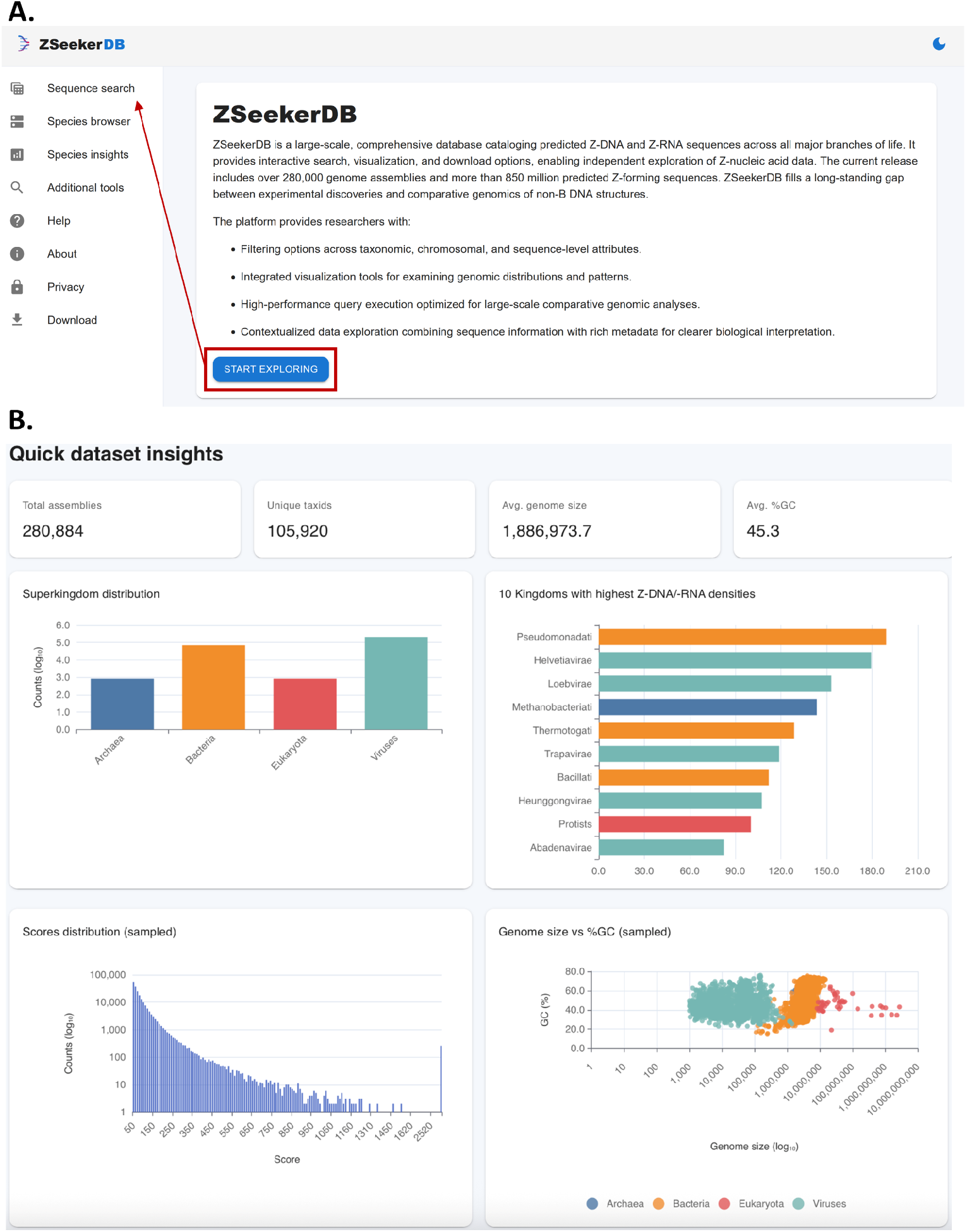
ZSeekerDB Homepage and navigation. **A**. Homepage and navigation tabs. **B**. Summary statistics and visualizations.

The homepage also features summary metrics for the entire dataset, as well as 4 summary plots (**Fig. 1B**). Specifically, the counts of assemblies in each superkingdom, the 10 kingdoms with the highest Z-DNA/-RNA densities, the distribution of predictions’ scores and the distribution of G+C content and genome size are displayed.

### Sequence search

The Sequence Search interface enables precise and flexible retrieval of candidate Z-nucleic acid sequences across organismal genomes. Queries can be constructed using a combination of parameters, including Species/Assembly (with autocomplete support), Chromosome, Genomic range (Start and End), Z-DNA Score (defined through numerical thresholds or range sliders), and “Sequence contains” (substring match) (**Fig. 2A-B**). Each search can be anchored to a specific assembly through URL parameters, ensuring reproducibility and deep linking across user sessions.

**Figure 2:**
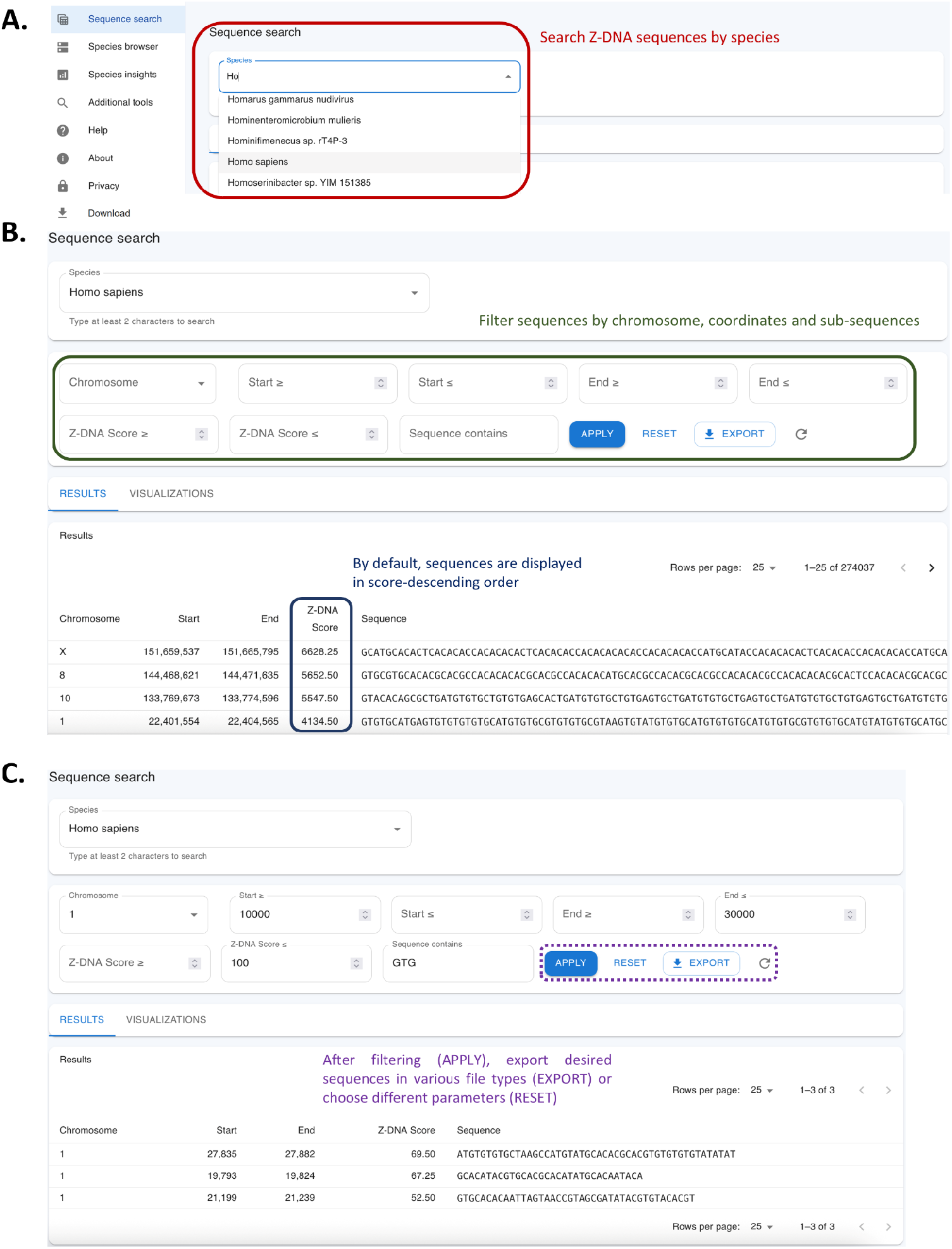
The sequence search interface page. **A**. The search panel shows species, chromosome, score and sequence filters. **B**. Results table with server-side pagination and column sorting. **C**. Export dialog demonstrating multiple formats.

Search results are presented in two complementary views, the Results and the Visualization tabs. This dual tabular and visual representation enables users to seamlessly transition between quantitative inspection and qualitative pattern recognition, enhancing both the interpretability and reproducibility of genome-wide Z-DNA analysis workflows (**Fig. 2B**). In the Results tab, retrieved loci are displayed in a virtualized, sortable data table with server-side pagination. Each entry includes its Chromosome, Start and End coordinates, Z-DNA Score, and nucleotide Sequence, allowing rapid inspection and comparative evaluation of individual candidates. This view supports filtering and direct data export in multiple structured formats, including CSV, TSV, JSON, XML, and Excel-compatible tables, using server-side streaming to accommodate large-scale downloads (**Fig. 2C**).

### Analysis and visualization pages

In the Visualizations tab, query data is represented graphically through a suite of dynamic plots. These include a histogram displaying the frequency distribution of predicted Z-DNA scores across all candidate loci within the selected dataset or assembly (**Fig. 3A**). Each bar represents the number of loci falling within a given score range, providing an overview of how Z-DNA propensity values are distributed genome-wide. Manually comparing histograms across species reveals shifts in the prevalence of strong Z-DNA propensity.

**Figure 3:**
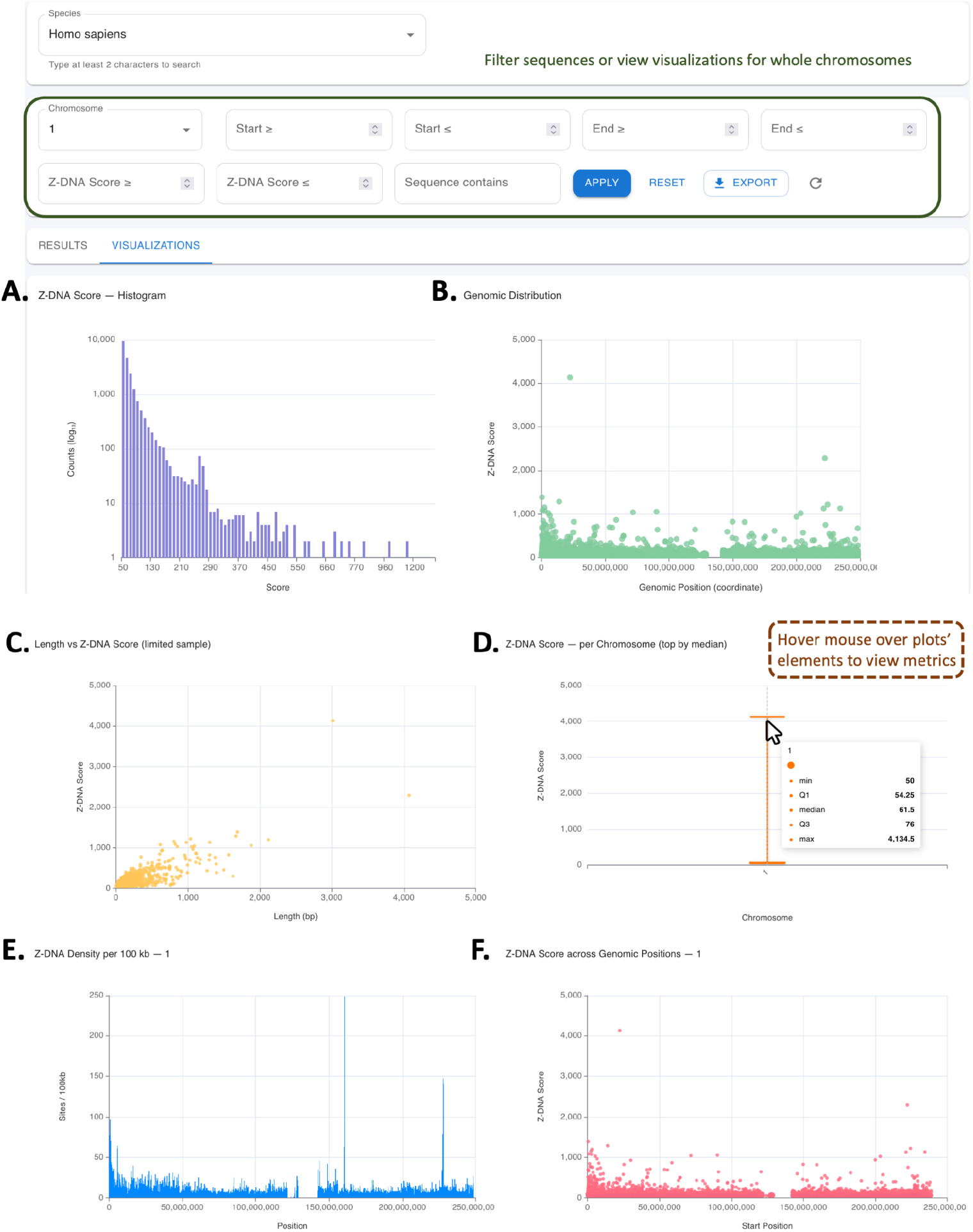
Sequence search visualizations. **A**. Histogram (with adjustable bin size) summarizes the distribution of Z-DNA scores for the current selection (species and genome region). **B**. Genomic distribution. **C**. Predictions’ length versus Z-DNA score scatter plot. **D**. Z-DNA score per chromosome (top 15 by median). **E**. Z-DNA locus density per 100 kb window illustrating the spatial uniformity of predicted sites. **F**. Genomic start coordinate versus Z-DNA score showing score dispersion and distribution patterns along the same chromosome.

For any selected species and chromosome combination, ZSeekerDB provides chromosome-specific visualization panels that enable the spatial and quantitative examination of predicted Z-DNA loci along the genomic coordinate space. The Genomic Distribution panel (**Fig. 3B**) displays the genomic positions of putative Z-nucleic acid sequences along the selected chromosome. This visualization provides an overview of whether the selected sequences are concentrated in particular genomic intervals or more evenly distributed across the chromosome, thereby highlighting potential clusters or hotspots of Z-nucleic acid propensity. The Length versus Z-DNA Score scatter plot (**Fig. 3C**) illustrates the relationship between the length of candidate Z-nucleic acid-forming sequences and their corresponding scores and a boxplot displays the range of scores across the selected chromosome (**Fig. 3D**). The “Density per 100 kb” histogram (**Fig. 3E**) represents the normalized distribution of Z-nucleic acid-forming sites along the selected chromosome, aggregated into fixed non-overlapping windows of 100 kilobases. Finally, the “Start versus Z-DNA Score” scatter plot (**Fig. 3F**) displays the genomic start coordinate of each candidate sequence on the x-axis and its corresponding Z-DNA score on the y-axis, for the specified chromosome. This visualization highlights whether high-scoring Z-nucleic acid sequences are clustered or dispersed and whether score magnitude correlates with particular chromosomal intervals.

Together, these chromosome-level visualizations complement the genome-wide summary plots by providing fine-scale spatial resolution, enabling researchers to identify candidate genomic regions that may warrant further investigation through comparative genomics and/or experimental validation.

### Species browser

The Species browser page enables faceted exploration of assemblies and taxa through fast distinct-value lookups and export functions (**Fig. 4A**). Filters are dynamically generated from the metadata schema and include categories such as Superkingdom, Kingdom, Phylum, Genus, Taxon name, Assembly level, and numeric ranges like Genome size or G+C content **(Fig. 4B**). Autocomplete widgets, powered by REST endpoints, return distinct values on demand to support efficient querying. The results are displayed in a main table listing assemblies or species that meet the selected criteria, with columns such as Taxon, Assembly, Level, Genome size, and G+C content. Users can easily export the filtered dataset through the /api/metadata/export endpoint. This page supports workflows such as “show all bacterial assemblies with G+C content between 35-45 and chromosome-level completeness” or “restrict to a particular genus”. The export capability allows the selected cohort to be used as an input list for subsequent sequence-level queries.

**Figure 4:**
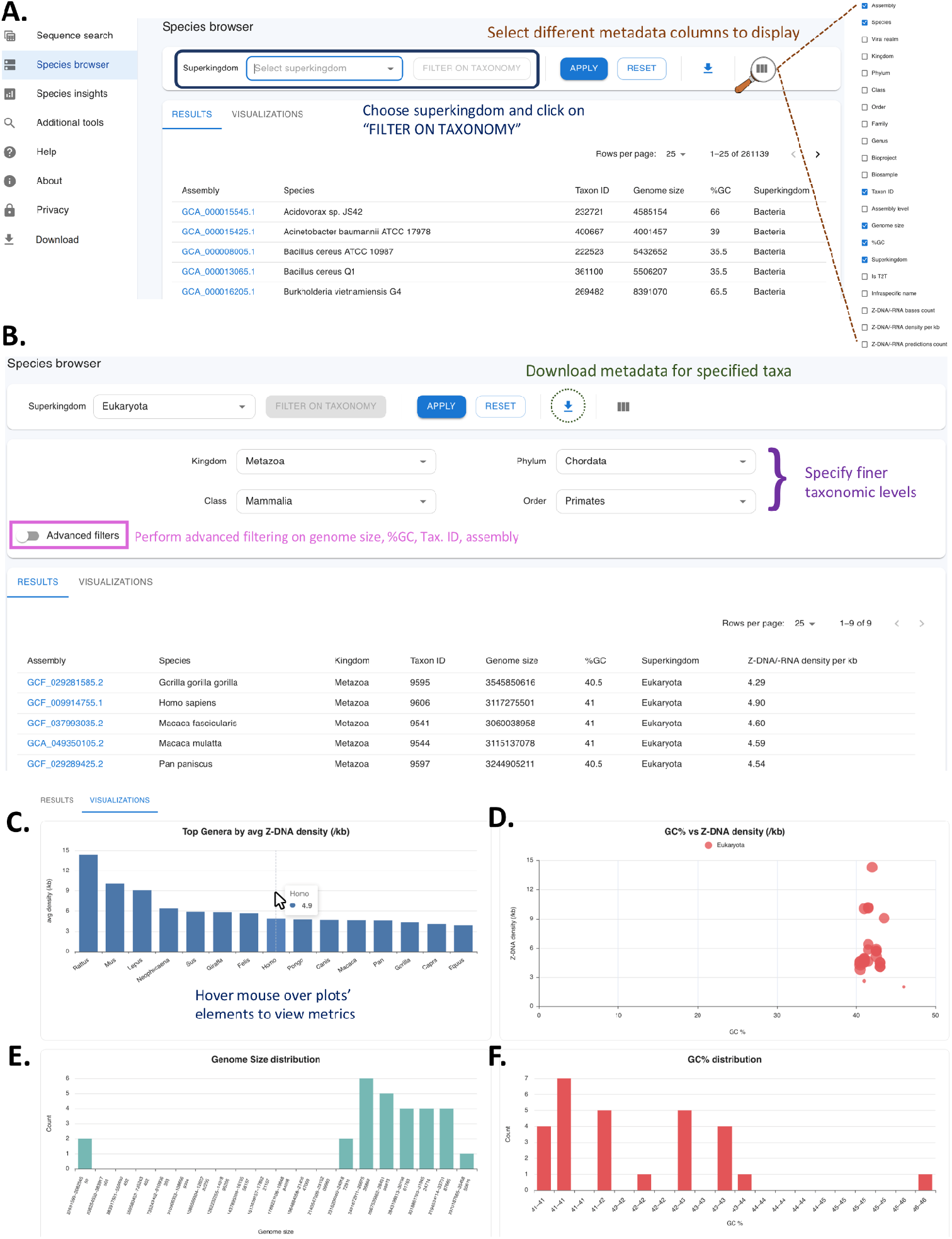
The Species browser user environment. **A**. Faceted metadata filters for taxonomic ranks and assembly attributes, including categories such as superkingdom, kingdom, phylum, genus, assembly level, genome size, and GC%. **B**. The corresponding list of assemblies with key descriptors: Assembly ID, Species name, taxonomic info, genome size, GC content and Z-DNA/Z-RNA density. **C**. Histogram of top genera ranked by average Z-DNA density. **D**. Scatter plot of GC% versus Z-DNA density per species. **E**. Histogram of genome size distribution for the filtered assemblies. **F**. Binned GC-content histogram across specified assemblies.

The Species Browser provides taxonomically resolved visual analytics of Z-DNA sequence density and genome-level features across the analyzed assemblies. Users can filter datasets by major taxonomic ranks (e.g., superkingdom, kingdom, phylum, class, order) and dynamically explore genome statistics and Z-nucleic acid sequence distribution patterns. The visualization panel summarizes these relationships through four complementary charts that collectively describe the structural and compositional variability of Z-nucleic acid formation potential among species (**Fig. 4C-F**).

The “Top Genera by average Z-DNA density” histogram (**Fig. 4C**), ranks genera according to their mean Z-nucleic acid sequence density, defined as the number of predicted Z-nucleic acid-forming base pairs divided by genome size. This bar chart highlights taxa enriched in Z-nucleic acid sequences, enabling comparative assessment of Z-nucleic acid-forming potential across evolutionary lineages and identification of genera exhibiting unusually high densities. The “G+C versus Z-DNA density” scatter plot (**Fig. 4D**) illustrates the relationship between genomic G+C content and Z-nucleic acid sequence density. The “Genome size distribution” histogram (**Fig. 4E**) summarizes the distribution of total genome sizes within the filtered taxonomic group. This view provides context for the genomic scale of the analyzed assemblies and allows users to assess how genome size variability may influence Z-nucleic acid sequence density normalization and comparative analyses. The “G+C distribution” histogram (**Fig. 4F**) depicts the frequency distribution of G+C content across genomes. Together, these four visualizations provide an integrative overview of genomic and compositional factors influencing Z-DNA sequence density, offering a comparative framework for exploring how taxonomic, structural, and sequence-level properties co-vary across the analyzed assemblies.

### Species insights

The Species Insights interface (**Fig. 5**) provides a taxonomically contextualized overview of the ZSeekerDB dataset, enabling users to explore large-scale patterns of Z-nucleic acid-forming sequence distribution across the tree of life. The page opens with a set of key statistics derived from the database metadata, including the total number of analyzed species and assemblies, as well as the ranges and mean values for genome size and G+C content (**Fig. 5A-B**). These summary metrics offer immediate contextual awareness regarding the taxonomic breadth and genomic diversity represented in the database. Taxonomic distributions are visualized through interactive bar and tree map charts that aggregate the number of assemblies and the total count of predicted Z-nucleic acid-forming sequences across hierarchical taxonomic ranks such as superkingdom, kingdom, phylum, and genus (**Fig. 5C**). The interface supports dynamic drill-down exploration, whereby user selections at higher taxonomic levels automatically update subordinate charts and filters, facilitating progressive navigation from broad phylogenetic groups to specific lineages.

**Figure 5:**
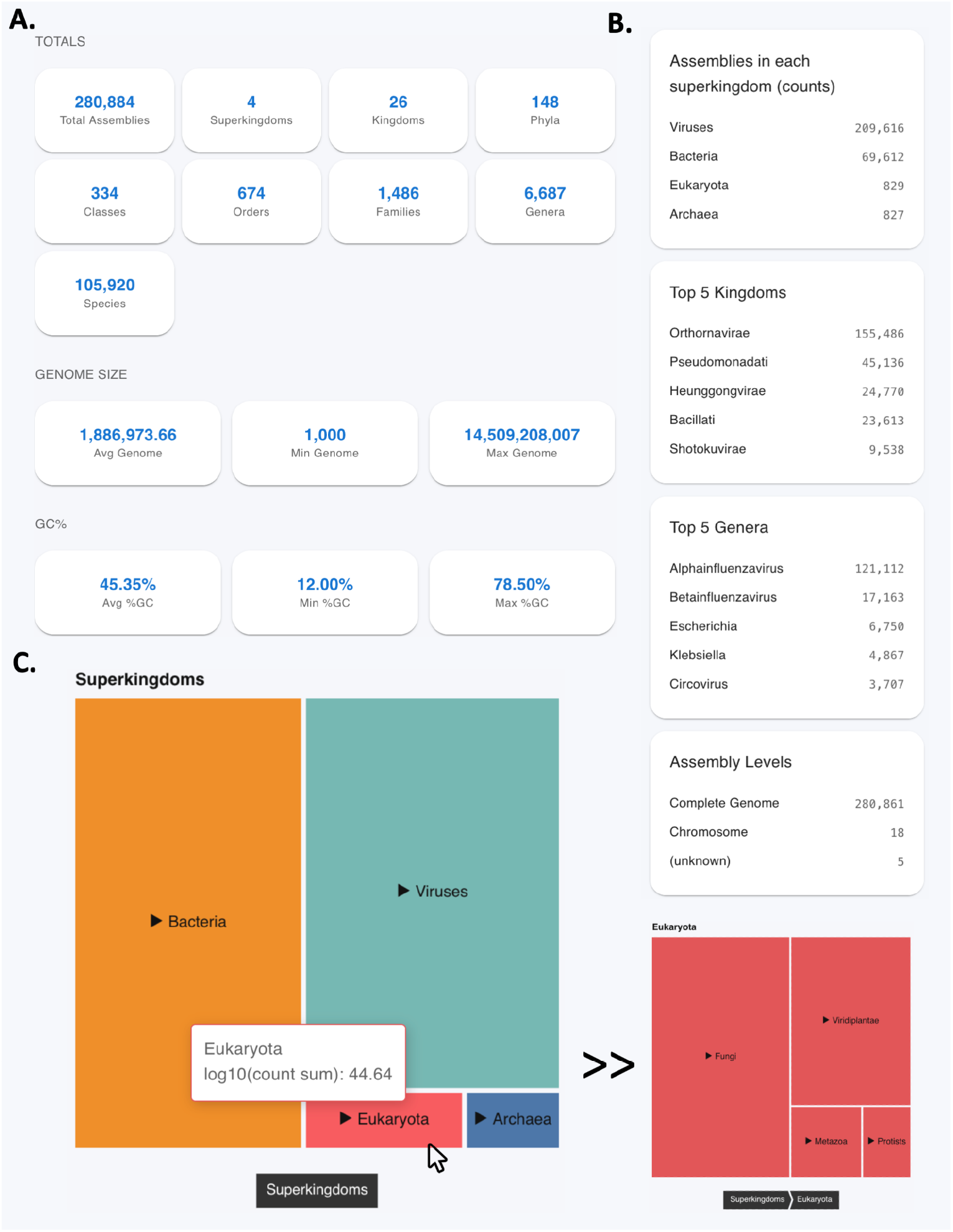
Species insights offer a quick view of the dataset attributes. **A**. Summary statistics and taxonomic distribution across superkingdoms. **B**. Drill-down by kingdom and genus. **C**. Tree map graph showing the proportion of assemblies in each taxonomic group, in log scale.

### Programmatic access

All visualizations are served through REST endpoints that return compact JSON outputs. For advanced analyses, a read-only SQL endpoint (/api/sql?query=…) allows parameterized SELECT queries, providing the same data used by the frontend. This design enables computational notebooks and pipelines to fully reproduce results shown in the interface. To ensure long-term stability and reproducibility, all API endpoints are versioned with each software release, and the complete database file is archived on Zenodo at https://zenodo.org/records/17533316.

## Discussion

Here, we introduce ZSeekerDB, the first database dedicated to Z-nucleic acid-forming sequences. Prior studies have addressed Z-nucleic acid-forming sequences at the level of individual genomes or limited taxonomic groups (10, 25, 30), but no study has provided a comprehensive resource spanning all major biological domains. The ZSeekerDB database encompasses Z-nucleic acid-forming sequence annotations for over 280,000 organismal genome assemblies, of organisms spanning the tree of life. Our website also provides an intuitive, interactive interface with search and filtering tools, dynamic queryable tables, visualizations, and downloadable data for independent analysis.

Multiple studies have characterized the diverse biological roles and genetic instability associated with Z-DNA and Z-RNA sequences (4, 13, 31, 32). These include their involvement in antiviral defense (12), activation of necroptosis pathways (33), regulation of gene expression (9, 10), and contributions to genome instability (2). ZSeekerDB provides a foundation for hypothesis generation regarding the evolutionary conservation, lineage-specific emergence and context-dependent functionality of Z-nucleic acid-forming sequences.

We anticipate that ZSeekerDB will be widely adopted by researchers seeking to investigate Z-nucleic acid-forming sequences across species. By enabling systematic exploration of Z-nucleic acid-forming sequences, ZSeekerDB will advance our understanding of their roles in organismal evolution, genome instability, and biological function.

## Supplementary data

Supplementary Table 1: Genomic assemblies and corresponding species of ZSeekerDB.

[ZSeekerDB Assemblies and Species](https://docs.google.com/spreadsheets/d/1XzOM6Ds9fZC7rinnu34diDJx8dcFUqraFguNOTKozSY/edit?usp=sharing)

## Code and data availability

ZSeekerDB is publicly available at https://zseeker-db.com/. The ZSeekerDB dataset can be found in Zenodo with a stable version https://zenodo.org/records/17533316. The GitHub code for the extraction of Z-DNA sequences, filtering and for the database and website are found in https://github.com/Georgakopoulos-Soares-lab/zdna.

